# Rapid evolution of sex role specialization in a hermaphrodite under sex-limited selection

**DOI:** 10.1101/2022.04.21.489077

**Authors:** Anna K. Nordén, Steven A. Ramm, Jessica K. Abbott

**Author notes:** Corresponding author. Biology Department, Lund University, Sweden. Author contributions: AKA, SAR, and JKA conceived the study. AKA carried out the data collection and analysis. AKA and JKA wrote the manuscript with input from SAR.

## Abstract

The evolution of separate sexes from hermaphroditism is thought to have occurred independently many times, and is linked to the evolution of sex chromosomes. Even though we have a good understanding of the theoretical steps in the evolution of sex chromosomes from a hermaphrodite ancestor, the initial stages are still hard to study because many sex chromosome systems are old. We addressed this problem by experimentally selecting a hermaphrodite via sex-limited experimental evolution for several generations, simulating the early stages in the evolution of a sex chromosome. More specifically, we used a GFP (green fluorescent protein) marker as a proxy for a sex-determining locus, and selected replicate populations of the simultaneously hermaphroditic flatworm *Macrostomum lignano* for fitness via the male sex role, female sex role, or both (i.e. a control). After 14 generations, a fitness assay revealed clear evidence for incipient sex role specialization, presumably reflecting the release from constraints usually imposed by selection on the other sex role. Importantly, however, this was not simply explained by differential sex allocation in the different selection regimes - insofar as morphological traits reflect the underlying trade-off over resource allocation to the male and female sex functions - because testis and ovary sizes did not diverge among treatments. Our study shows that sex role specialization can occur rapidly as a result of sex-limited selection, which is consistent with genetic constraints between sex-roles, and in line with the first predicted steps towards the evolution of a new sex chromosome system.

## Introduction

Most animal species have separate sexes, but about 5% are hermaphroditic (Jarne and Auld 2006). Within animals, it is thought that transitions from hermaphroditism to separate sexes (known as dioecy in plants and gonochorism in animals) have occurred several times independently (Avise 2011). There are several possible evolutionary scenarios for such transitions; at one extreme is a slow scenario with a gradual increase in investment in one or the other sex role eventually leading to the evolution of separate sexes, and at the other is a rapid scenario with fixation of mutation(s) causing sterility in one or both sex roles with subsequent specialization (Bachtrog et al. 2014). A combination of slow and rapid processes is also possible, for example when an increase in the frequency of male-sterility alleles results in the production of a gynodiecious system (i.e. of females plus hermaphrodites), which then causes gradual specialization of hermaphrodite individuals in the male sex role (Avise 2011). When sexual differentiation is genetically determined, then an evolutionary transition to separate sexes will result in the evolution of sex chromosomes, which in many separate-sexed species are ultimately responsible for most aspects of sexual dimorphism (Wei and Barbash 2015).

The theory of evolutionary transitions from hermaphroditism to separate sexes is well-established, and the standard scenario starts with the establishment of a sex-determining region on an autosome (Charlesworth and Charlesworth 1978, Bachtrog et al. 2014, Beukeboom and Perrin 2014). A minimum of two mutations is needed to fully transition to separate sexes: one mutation generating male-sterility in females and one generating female-sterility in males, otherwise a mixed mating system results (i.e. gynodioecy or androdioecy; Charlesworth and Charlesworth 1978, Charlesworth et al. 2005, Perrin 2009, Beukeboom and Perrin 2014). Recombination cessation later evolves around the sex determining region, thereby preventing the production of sterile individuals. The region of recombination cessation can then increase to encompass other genes with sex-specific effects which are located close to the sex determining region (Beukeboom and Perrin 2014). Eventually, as the non-recombining region increases and degenerates, highly heteromorphic sex chromosomes can evolve (Bachtrog et al. 2011).

The fixation of a female- or male-sterility mutation in a population of hermaphrodites, initiating a potential transition to separate sexes, is likely to usually be associated with some sort of selective advantage. The most plausible scenario involves trade-offs between male and female fitness in hermaphrodites. Since hermaphrodites need to invest energy into both reproductive systems, they may be less efficient in producing gametes for a given resource investment compared to separate-sexed individuals (Charnov et al. 1976). This can lead to conflict between sex roles, either over resource allocation to each sex function (Heath 1977), or else because it is developmentally or physiologically difficult to be equally efficient at producing different types of gametes (Abbott 2011). A resolution to this conflict therefore could be to specialize in one sex function to increase reproductive output, providing scope for an initial selective advantage to a sex-specific sterility mutation within a population of hermaphrodites (Bachtrog et al. 2014). There is considerable literature investigating potential trade-offs between sex roles in hermaphrodites. Although unambiguous evidence has proven challenging to obtain, likely due to differences between individuals in overall resource acquisition and/or trade-offs with somatic investment, several studies have provided unequivocal evidence of the existence of trade-offs between male and female function (Schärer 2009, Schärer and Pen 2013).

Empirical observations of young sex chromosome systems are generally consistent with the theory outlined above. For example, work in the gynodioecious wild strawberry (*Fragaria vesca* ssp.) has demonstrated that female plants have a dominant male sterility gene at a single location on chromosome 4, suppressing the male function (Tennessen et al. 2013). Similarly, the dioecious herb *Mercurialis annua* has a male determining region that has lower recombination rates than expected, with male-specific genes located around the region (Veltsos et al. 2019). Zemp et al. (2018) also found clear evidence of sexual specialization in *Silene latifolia* within 5 million years after the evolution of separate sexes in this species. Nevertheless, gaps remain in our understanding of the early stages of sex chromosome evolution from a hermaphroditic ancestor, since most of the best-studied sex chromosome systems are ancient, and even comparatively young sex chromosome systems may often be several million years old (Bachtrog et al. 2011). Outstanding questions include what kinds of mutations are usually involved in the transition to separate sexes, how quickly sex-specific specialization can evolve, and the extent to which the properties of the sex-linked regions evolve in response to new selective situations caused by the presence of the sex-determining locus (Charlesworth 2019). It is also unclear how trade-offs between sex roles may play into sex-specific specialization during early sex chromosome evolution, although theory suggests that loci under sexually antagonistic selection for fitness (potentially associated with different allocation strategies) are likely to play an important role (Olito and Connallon 2019).

To gain a better understanding of transitions from hermaphroditism to separate sexes in real time, we subjected the simultaneously hermaphroditic flatworm *Macrostomum lignano* to sex-limited experimental evolution, simulating the evolution of a novel sex chromosome. We used a genetic marker (green fluorescent protein, GFP) as a “sex-determining gene”, i.e. meaning that individuals carrying the marker could only gain fitness through either their male or female sex function, not both. We subjected 4 replicate populations within each selection regime to either male-limited selection (fitness through the male sex role), female-limited selection (fitness through the female sex role), or to a control treatment. In this set-up the marker acts via our experimental protocol as a proxy for a dominant sterility mutation in one sex function, where “sterility” is enforced via the selection protocol (but does not cause true sterility). Hence, we simulate the formation of a nascent male-dominant XY-system in the male-limited selection regime and a nascent female-dominant ZW-system in the female-limited selection regime. After 15 generations we investigated the response to selection in sex-specific fitness compared to control populations. Additionally, we looked for evidence of morphological changes by measuring the overall body size of worms and the relative sizes of testes and ovaries after 25 generations of selection.

We expected that populations would respond to the selection regimes by increasing their fitness in the direction of selection, i. e. that the male-selected worms would increase their fitness in the male sex role, and the female-selected worms would increase their fitness in the female sex role. Additionally, if there are important trade-offs between sex roles, any increase in fitness in one sex role fitness should result in a decrease in fitness in the other sex role. We also predicted that changes in sex-specific fitness would be accompanied by morphological changes, for example that male-selected populations should increase their (absolute or relative) investment in testes.

## Methods

### Study species

*Macrostomum lignano* (Macrostomorpha, Platyhelminthes) is a small, free-living marine flatworm found in the intertidal zones of beaches around the Adriatic Sea in Northern Italy (Ladurner et al. 2005, Wudarski et al. 2020). It primarily feeds on diatoms and is transparent, which is ideal when measuring the size of organs such as ovaries and testes (Schärer and Ladurner 2003). *M. lignano* has paired ovaries and testes located along each side of the gut and mates reciprocally, meaning that both partners donate sperm to fertilize the eggs of their partner (Schärer et al. 2004, Ladurner et al. 2005, Vizoso et al. 2010). Despite being a simultaneous hermaphrodite, *M. lignano* never self-fertilizes (Vizoso et al. 2010).

In the laboratory, worms are kept in populations of 100 individuals in glass petri dishes, with f/2 medium (Andersen et al 2005) and fed *ad libitum* with the diatom algae *Nitzschia curvilineata* in a constant environment of 20 °C and 60% humidity on a 12:12 h light:dark cycle. Laboratory lines used for the creation of start-up populations for the experimental evolution in this study were the transgenic GFP(+) line BAS1, and wild-type population LS2. BAS1 is homozygous for the GFP gene. Detailed information about the creation and maintenance of these populations can be found elsewhere (Marie-Orleach et al. 2014, Zadesenets et al. 2016). The GFP gene is dominant and ubiquitously expressed, and GFP(+) worms fluoresce under a near-UV light source. The marker has previously been shown not to affect morphology, mating success, or mating rate compared to wild-type lines (Marie-Orleach et al. 2014). In order to obtain genetically variable start-up populations, we crossed homozygous BAS1 worms with LS2 wild-type worms, which produced worms heterozygous for the GFP marker (+/-). Next, we backcrossed these worms with LS2, creating ~50% heterozygous offspring and ~50% unmarked (GFP(-)) offspring. These constituted the start-up populations; the heterozygous GFP(+) offspring were used for the male/female/control treatments and the GFP(-) individuals were allocated to the “source” populations (see below).

### Experimental evolution protocol

The experimental evolution lines consist of four replicate populations (denoted 1-4) within each selection regime (female-limited, male-limited and control), resulting in 12 populations in total (denoted F1-4, M1-4 and C1-4). Populations with the same replicate number are not more closely genetically related to each other than to other replicates, but they are related in terms of handling (i.e. culturing was during the same time of the day, and worms with the same replicate number were placed in the same area of the incubator).

A new generation starts with 48 mature, GFP(+) individuals from each replicate population being crossed with two worms each from the matched “source” population. Source populations are maintained in the same way as the other laboratory lines described above, and have the same genetic origin as the GFP(+) individuals from the same combination of treatment and replicate. They are used to provide mating partners for the GFP(+) individuals that neither carry the marker nor are exposed to the sex-limited selection. The trios of worms (1 GFP(+) and 2 GFP(-) individuals) are held in individual wells of 24-well tissue culture plates for one week, to provide opportunities for sperm competition and mate choice (figure 1). Therefore, the effective population size for each experimental line was approximately N_e_ ≈ N + 0.5 = 144.5 (i.e. 48 GFP(+) individuals and 96 GFP(-) individuals; Caballero 1994, Falconer and Mackay 1996), which is similar to several previous experimental evolution studies (e.g. Prasad et al. 2007, Michalczyk et al. 2011, Innocenti et al. 2014, Buechel et al. 2016). Only 48 individuals per population experience sex-specific selection each generation, but we expected that this selection would be sufficient, given the success of other experimental evolution studies similar to our design (e.g. Morrow et al. 2008).

**Figure 1:**
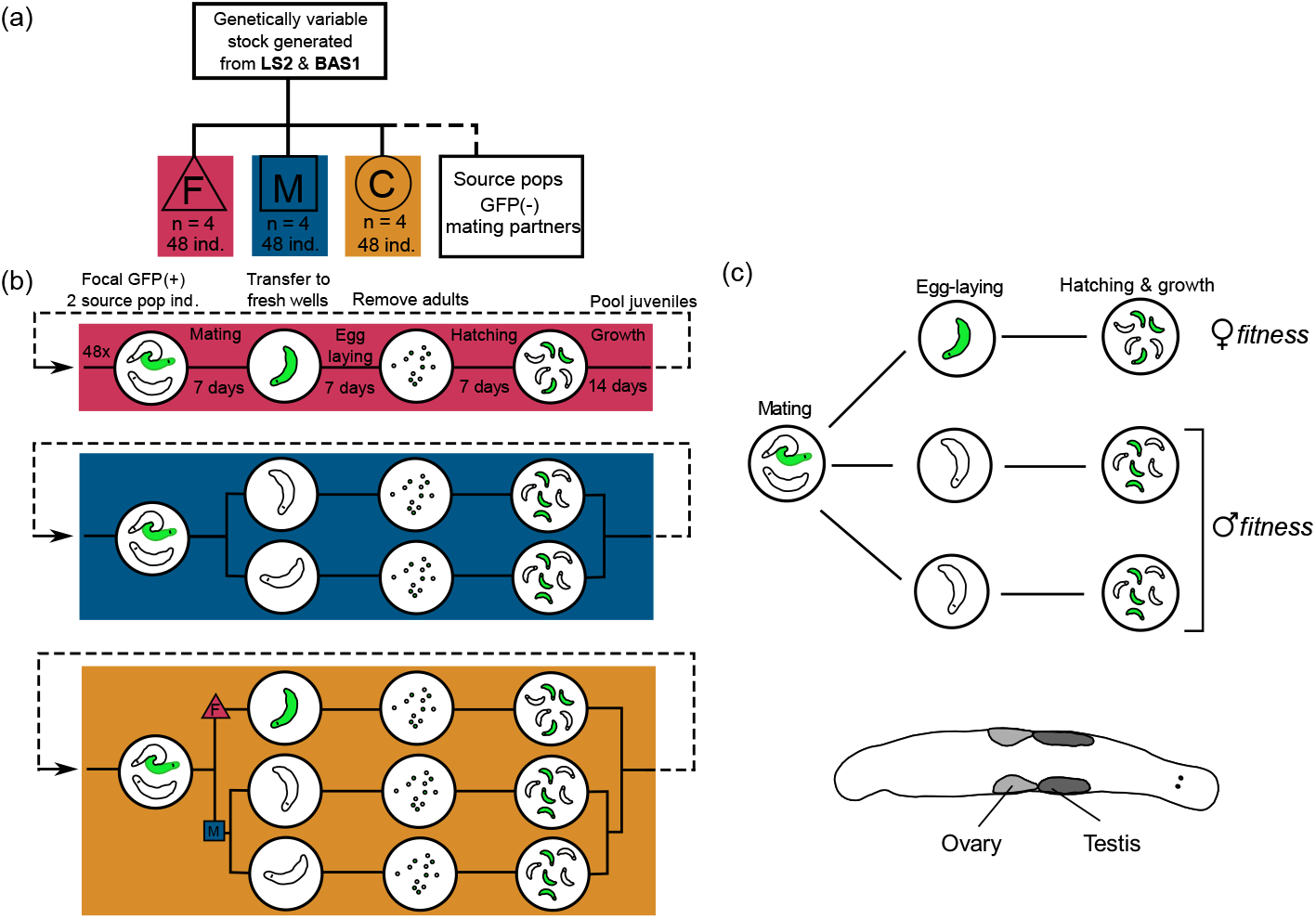
An overview of the selection regimes (a), the sex-limited selection protocol (b), and the measures of the response to selection (fitness assay and morphological measurements; c).

After 7 days for mating, worms are isolated in new wells to lay eggs for one week. In the female-limited selection regime, the GFP(+) focal worms are isolated so that the GFP-marker is inherited via the female sex role, i.e. eggs. In the male-limited selection regime the two GFP(-) mating partners are isolated for egg laying, so that the GFP-marker is inherited via the male sex role, i.e. sperm. Finally, in the control lines, half of the selected worms are treated in the same way as the female-limited selection regime, and half are treated in the same way as the male-limited selection regime. Egg-laying worms are discarded after one week, and eggs are left to hatch in the wells. After one week of growth, offspring in each well are moved to petri dishes to mature. If an experimental line does not produce sufficient numbers of GFP(+) offspring, back-up offspring from the previous generation are used, so that generations are mainly non-overlapping, but not completely so.

During the first ten generations, each generation lasted for four weeks, but due to poor production of offspring within the female selection regime, we subsequently extended the maturation period to two weeks instead of one, which was successful in increasing offspring numbers. We based this decision on prior knowledge that in juveniles, testes mature slightly earlier than ovaries, so that female-limited selection could have resulted in longer ovary maturation times (Vizoso and Schärer 2007).

### Fitness assay

Worms from the 14^th^ experimental generation were collected and isolated in wells, directly after the completion of egg-laying to produce generation 15. More specifically, mating partners not used in the egg-laying were held in new wells instead of being discarded, and after egg laying for generation 15 was complete, egg-laying worms were combined with the same partners again to provide 24 trios per selection regime and replicate. These trios of worms were again allowed to interact for 7 days. Next, all individuals were isolated in new wells to lay eggs for 7 days. Adult worms were then discarded, eggs were left to hatch, and juveniles to grow. The whole procedure (mating for 7 days followed by egg laying for 7 days, and growth of juveniles) was then repeated using the same trios of individuals, in order to increase total offspring production and thereby decrease the error in the individual fitness measurements. Fitness was measured as number of GFP(+) and GFP(-) offspring per well (i.e. per GFP(+) focal individual). Total number of offspring produced via eggs was used as a measure of female fitness, and proportion GFP(+) offspring produced by both GFP(-) mating partners was used as a measure of male fitness. This fitness assay builds on a standard fitness assay protocol commonly used in *Drosophila* (e.g. Abbott et al. 2013, Lund-Hansen et al. 2020)and is essentially the same as the sex-limited selection protocol, except for the fact that all worms are given the opportunity to lay eggs. We measured sex-specific fitness for 267 individuals in total (91 from the control regime, 91 from the male-limited regime and 85 from the female-limited regime).

### Phenotypic measurements

Body size, relative testis size and relative ovary size were estimated with a standard method used in this species (Schärer and Ladurner 2003). Briefly, worms were first isolated in wells containing f/2 solution and starved overnight. Each worm was then anesthetized for ten minutes in a well containing a mix of 600 μl artificial seawater (ASW) and 1 ml MgCl2 –solution (conc 7.14 mg/ml). It was then slightly squeezed between a microscope slide and a cover slip, with pieces of plastic film of a standard thickness used as spacers in between (Schärer and Ladurner 2003). A picture of the whole body was then taken at x40 magnification, and of the ovaries and testes at x200 magnification. Photos were processed and analyzed in the program ImageJ (version 1.51), where the area of the body and area of the ovaries (sum of left and right) and the testes (sum of left and right) were calculated, respectively. Worms were photographed in random order, and the observer was blind with respect to the selection regime. We determined the repeatability of the morphological measurements by measuring 20 pictures three times for testes size, ovary size and body area.

This assay was carried out at generation 25. After discarding poor-quality images, our final sample size was 121 for body area (40 from the control regime, 40 from the male-limited regime, and 41 from the female-limited regime), 116 for testes area (39 from the control regime, 37 from the male-limited regime, and 40 from the female-limited regime), and 115 for ovary area (40 from the control regime, 36 from the male-limited regime, and 39 from the female-limited regime). The repeatability of the morphological measurements was high (body: intraclass correlation coefficient, ICC = 0.998, p = < 0.001, testes: ICC = 0.940, p = < 0.001 and ovary: ICC = 0.882, p= < 0.001; Koo and Li, 2016, Vaz et al 2013).

### Statistical analysis

All statistical analysis was carried out in R Version 4.1.0 (R Core Team 2021). For the fitness assay, we excluded data from worms that had no offspring in either sex role (which could for example occur if the worm was injured during handling) and individuals that had undefined male fitness (i.e. no offspring from either mating partner at all). The sex-specific fitness measures were standardized to a mean of zero and standard deviation of one to make them directly comparable. Changes in sex-specific fitness between selection regimes were analysed using a mixed model approach implemented in lme4 (Bates et al. 2015) with selection regime, sex role (male or female), and their interaction as fixed effects, and individual ID and replicate population nested within selection regime and sex role as random effects (Lund-Hansen et al. 2020, Manat et al. 2021). Since we have two types of sex-specific fitness measures per individual, the random effect of ID was to control for variance arising from overall fitness differences between individuals, and the nested effect of replicate population was included in order to avoid pseudoreplication (Arnqvist 2020). Posthoc comparisons were carried out using emmeans (Lenth 2021).

Because gonad size is correlated with overall body size, we used relative gonad areas as a measure of investment in testes and ovaries (i.e relative testes area was calculated as testes area/body area, and relative ovary area was calculated as ovary area/body area). Differences in body area, relative testes area, and relative ovary area were each analysed with mixed models where selection regime was a fixed effect, and replicate population nested within selection regime was a random effect. Although here we report results from analyses using relative gonad sizes, results were qualitatively similar if body area was instead included as a covariate (see Results).

To examine whether there was a trade-off between investment in testes and ovaries, and if the magnitude of this trade-off differed between selection regimes, we first calculated the correlation between relative testes area and relative ovary area within each combination of selection regime and replicate population. We then used these 12 correlation coefficients as the dependent variable in a one-way anova analysis, with selection regime as the independent variable. The aim of this analysis was to test for consistent differences in the correlation coefficient between selection regimes. Since there was no evidence of any differences between treatments (see Results), we also tested if the mean correlation coefficient across all replicate populations was significantly different from zero using a one-sample t-test.

Finally, we tested whether differences in morphology could explain differences in sex-specific fitness. Since morphological data and sex-specific fitness data were collected at different times, this analysis could not be carried out on the individual level, so instead we used replicate population mean values. We used a regression approach with sex-specific fitness as the dependent variable, and morphological variable (body area, relative testes area, or ovary area) as the predictor variable. The effect of selection regime was not included since replication was so limited in this analysis. However, results were qualitatively similar when carrying out an ancova analysis including both the morphological variable and selection regime as predictors (data not shown).

## Results

### Fitness assay

Consistent with our main hypothesis that increased fitness in one sex role should come at a cost to the other sex role, there was a significant interaction effect between treatment and sex role on fitness (Table 1, Figure 2). Posthoc testing was restricted to comparisons within each sex role, in order to minimize the need for correcting for multiple comparisons. We found that there was a trend towards a significant difference between the F and C regimes in the female sex role (p=0.07), and between the F and M regimes in the male sex role (p=0.08). From Figure 2, it is clear that the female-limited selection regime has higher fitness through eggs and lower fitness through sperm compared to the other two selection regimes.

**Table 1:** Results of the mixed model analysis of sex-specific fitness.

| Effect | MS | Num<br>DF | Den DF | F-value | P-value | Variance |
| --- | --- | --- | --- | --- | --- | --- |
| **Selection regime** | 0.1949 | 2 | 17.23 | 0.1994 | 0.8211 | - |
| **Sex role** | 0.0101 | 1 | 16.67 | 0.0103 | 0.9202 | - |
| **Selection\*sex role** | 4.9796 | 2 | 16.65 | 5.094 | 0.0188 | - |
| **ID** | - | - | - | - | - | 0.0137 |
| **Replicate** | - | - | - | - | - | 0.0004 |
| **Residual** | - | - | - | - | - | 0.9775 |

**Figure 2:**
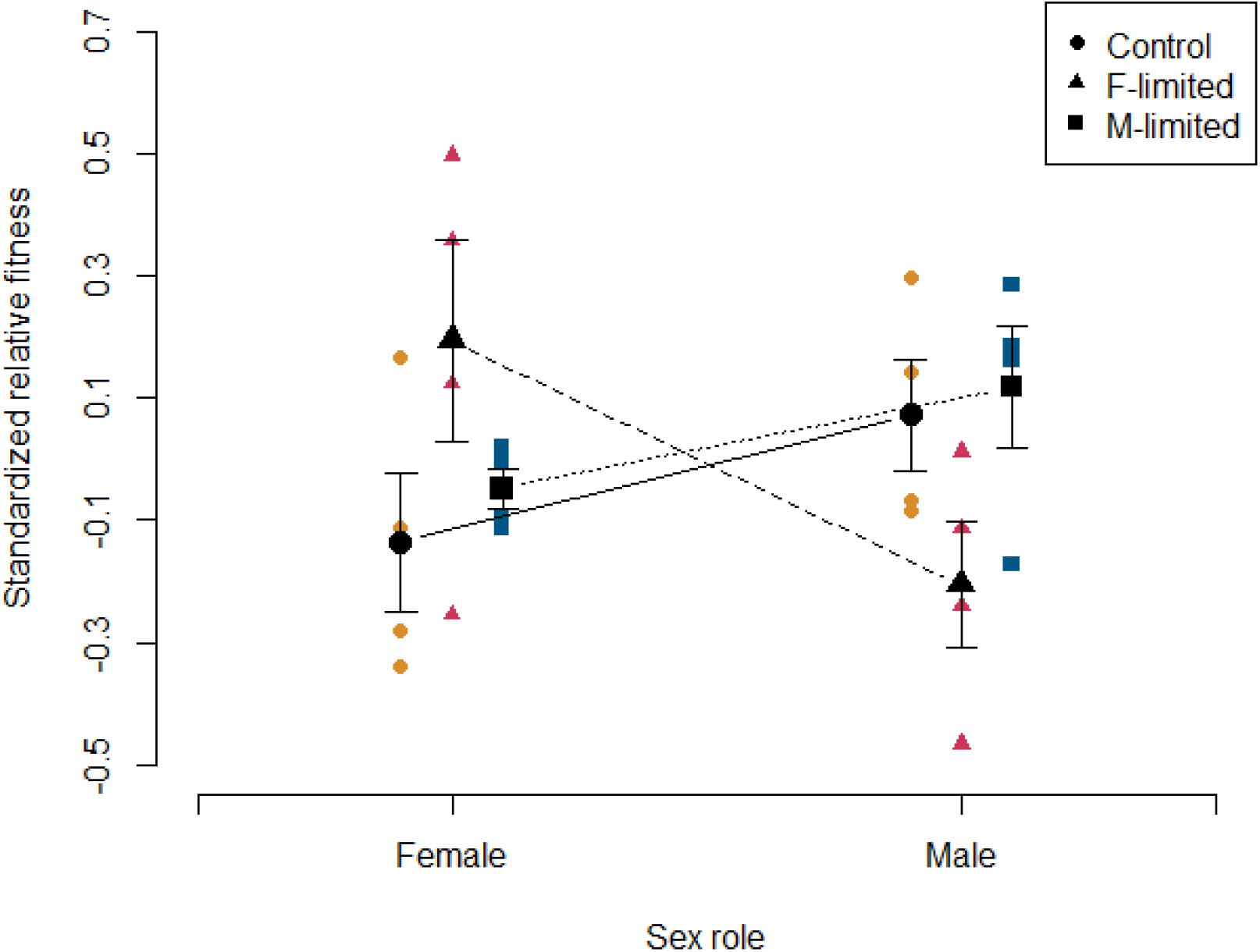
Interaction plot of standardized relative fitness dependent on sex role (male or female) and selection regime (female-limited, male-limited, or control). Small coloured points represent replicate population means.

### Phenotypic measurements

There were no significant differences between selection regimes for any of the morphological measurements (body area, relative testes area, and relative ovary area; all p>0.6), although body area was, as expected, significantly related to absolute gonad sizes (see tables S1-S3). This suggests that morphological differences are unlikely to explain differences in sex-specific fitness between selection regimes.

There was no evidence of a trade-off in investment in testes and ovaries since the mean correlation coefficient between relative testes size and relative ovary size across all replicate populations was positive, and significantly different from zero (one-sample t-test: *t* = 3.85 df = 11, p-value = 0.002717, mean = 0.314, 95% confidence interval = 0.134, 0.494). Nor was there any evidence that the correlation coefficient differed between treatments (one-way anova: *F_2,9_* = 0.836, p=0.464). Collectively, these results suggest that individuals from all selection regimes mainly differ in their total investment in reproduction, rather than trading-off investment in testes with investment in ovaries.

When testing whether morphological differences could explain differences in sex-specific fitness, the only robust result was for relative testes area, which was positively related to fitness in the male role (Table 2 and Figure 3A). However, there was also a trend that larger ovary area resulted in lower fitness in the female role (Table 2 and Figure 3B). This suggests again that differences in sex-specific fitness between the selection regimes cannot be explained by morphological differences. It is also consistent with previous studies which have used testes size as a proxy for investment in the male sex role (Schärer and Ladurner 2003, Vizoso and Schärer 2007, Janicke et al. 2013, Vellnow et al. 2017), but that ovary area is a poor predictor of fitness through the female sex role (Janicke et al. 2011).

**Figure 3:**
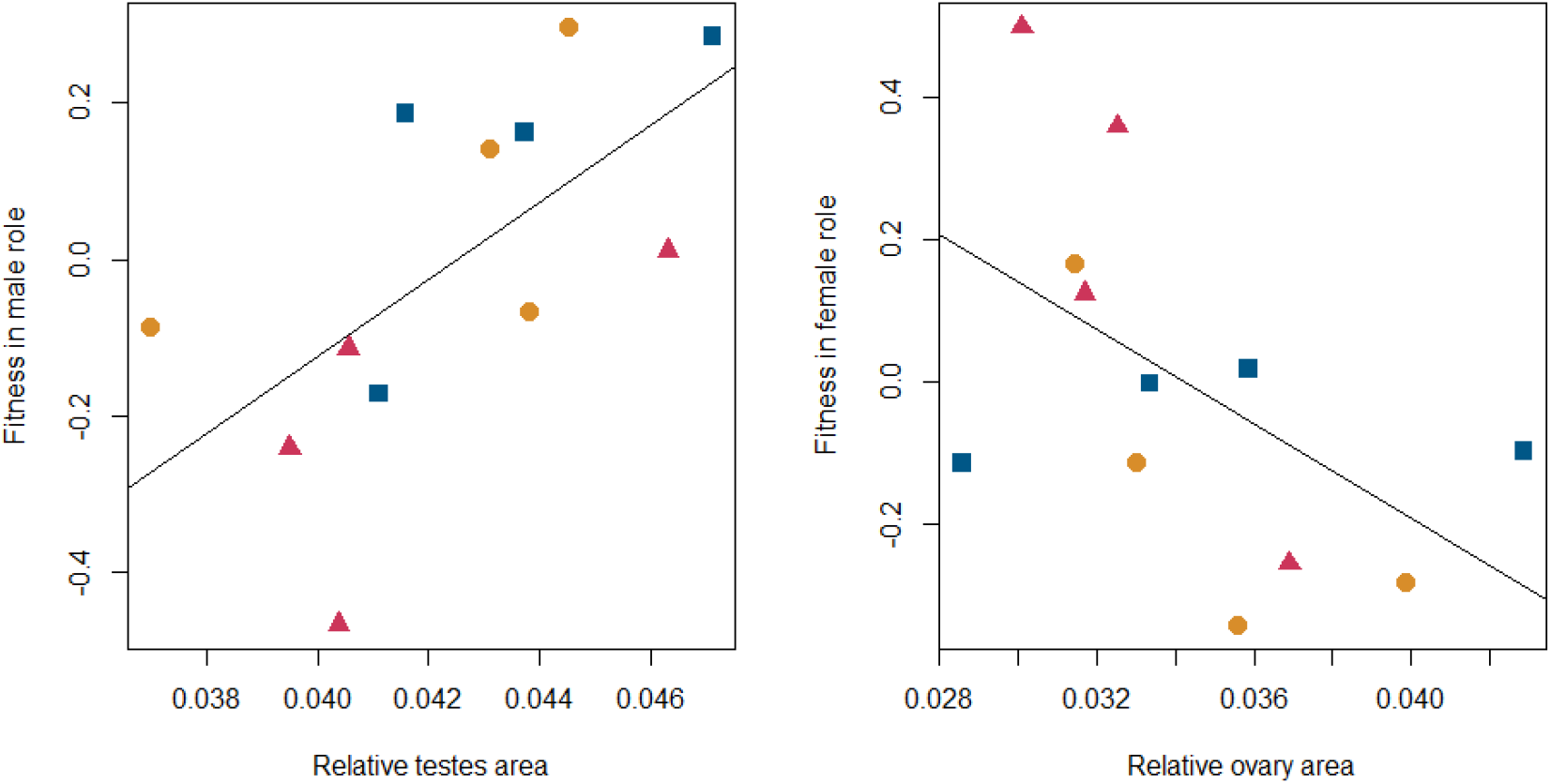
Scatterplots showing the relationship between sex-specific fitness and relative testes area (A), and relative ovary area (B). The symbols represent each selection regime (triangles for the female-limited regime, squares for the male-limited regime, and circles for the control regime).

**Table 2:** Results of the ancova analyses of the relationship between morphology and sex-specific fitness.

| Sex role | Effect | MS | Num DF | Den DF | F-value | P-value |
| --- | --- | --- | --- | --- | --- | --- |
| **Female** | Body area | 0.0626 | 1 | 10 | 0.9578 | 0.3508 |
|  | Relative testes | 0.0512 | 1 | 10 | 0.7694 | 0.4010 |
|  | Relative ovary | 0.2060 | 1 | 10 | 4.0336 | 0.0724 |
| **Male** | Body area | 0.1256 | 1 | 10 | 2.7614 | 0.1275 |
|  | Relative testes | 0.2279 | 1 | 10 | 6.4634 | 0.0293 |
|  | Relative ovary | 0.1150 | 1 | 10 | 2.4698 | 0.1471 |

## Discussion

This is, to our knowledge, the first time a completely sex-limited selection experiment has been carried out in a simultaneous hermaphrodite (but see Bonel et al. 2018 for an example of experimental evolution via relaxed selection in the male or female sex role). Our aim with this experiment was to try to recreate in the laboratory the early stages of the evolution of a novel sex-determining system from a hermaphroditic ancestor. We expected to see evidence of sexual specialization as a result of our experimental protocol, consistent with predictions about early sex chromosome evolution (Charlesworth and Charlesworth 1978, Bachtrog et al. 2014, Beukeboom and Perrin 2014), and this expectation was generally fulfilled. We found evidence of a clear response to female-limited selection in fitness, such that female-selected lines had higher production of eggs but reduced performance in sperm competition compared to control and male-selected lines. These differences could not be directly attributed to changes in gonad size, since there was no significant difference between selection regimes in body area, relative testes area, or relative ovary area. However, mean testes area per line was positively related to mean male-specific fitness per line, suggesting that testes area may influence siring success, for example via increased sperm production (Marie-Orleach et al. 2016). For female fitness, other factors such as egg-laying rate may be of greater importance (Janicke et al. 2011).

### Rapid evolution of female-specific fitness

Consistent with results from gonochorists (e.g. Rice 1998, Prasad et al. 2007, Morrow et al. 2008, Immonen et al. 2014, Stångberg et al. 2020) and other simultaneous hermaphrodites (Janicke et al. 2016, Bonel et al. 2018), we found that sex-specific fitness evolved rapidly in response to altered sex-specific selection pressures. There are three potential mechanisms which could account for the increase in fitness via the female sex role and decrease in fitness via the male sex role in the female selection regime. These mechanisms are mutation accumulation in the unselected sex role, energetic trade-offs in investment between sex roles, or sexual antagonism between sex roles.

Our experimental set-up seems to have been successful in selecting for increased fitness via the female sex role, and the decreased in fitness in the male sex role could be accounted for by accumulation of male-deleterious mutations. Results from Bonel et al. (2018) provide some support for this interpretation. They carried out experimental evolution with relaxed selection via either the male or female sex role, and found that relaxed selection in the male sex role resulted in reduced juvenile survival in the simultaneously hermaphroditic snail *Physa acuta.* They interpreted this finding as resulting from reduced purifying selection via the male sex role, suggesting that male-specific fitness may be particularly sensitive to mutation accumulation. Nevertheless, Bonel et al. (2018) did not detect any differences in adult sex-specific fitness in their study, in contrast to results presented here.

A second possibility is that the female-selected lines have altered their sex allocation. As discussed in the introduction, theories of sex allocation in hermaphrodites suggest that fitness via one sex role should come at a cost to the other (Heath 1977, Charnov 1982, Schärer 2009). In our experimental set-up, these theories would suggest that female-selected lines have evolved to invest more in egg production at the expense of sperm production. Although this prediction is consistent with our results, changes in sex-specific fitness are unlikely to be mediated by changes in ovary size, since relative ovary area showed a tendency to be negatively related to fitness via the female sex role (figure 3B). Nevertheless, a handful of studies have successfully demonstrated trade-offs between male and female sex function in this and other species (Schärer 2009, Di Bona et al. 2015, Picchi and Lorenzi 2019, Brand et al. 2022), so perhaps ovary area does not capture the most important aspects of investment in the female sex role. This is in line with results from Janicke et al. (2011), which reported that neither body size nor ovary size was correlated with female fitness in *M. lignano*.

Finally, the changes observed in the female-selected lines could be explained by sexual antagonism, i.e. when genetic variants that increase fitness in one sex role but decrease it in the other. Sexual antagonism is well-studied in separate-sexed species (e.g. Cox and Calsbeek 2009, Barson et al. 2015, Collet et al. 2016, Dutoit et al. 2018, Ruzicka et al. 2019), but still not well-understood in hermaphrodites (but see Olito 2016, Olito et al. 2017, Olito et al. 2018, Olito and Connallon 2019 for theoretical treatments of this problem), and can be considered a form of antagonistic pleiotropy in these species (Abbott 2011, Schärer et al. 2014). Although a mutation which influences sex allocation could be classed as a form of sexual antagonism (e.g. if it increases investment in the female sex role at the expense of the male sex role), other types of sexual antagonism are possible in hermaphrodites - for example a mutation which increases efficiency of sperm production could pleiotropically reduce egg production without necessarily altering energetic investment in each sex role (Abbott 2011). Our results could therefore also be explained by an increase in the frequency of alleles which promote egg production but pleiotropically decrease sperm quality in the female-selected lines.

These three explanations are not mutually exclusive, and we cannot definitively distinguish between them with the data at hand. That being said, we do not feel that mutation accumulation in the male sex role is the most likely explanation since we measured fitness after only 14 generations of selection, making it rather unlikely that extensive mutation accumulation had occurred (Bonel et al. 2018 did not detect a significant effect of their selection regime until after 35 generations, for example). The sex allocation and sexual antagonism hypotheses are not easily distinguished, but the energetic trade-offs associated with the sex allocation hypothesis are perhaps unlikely to be an important selective constraint in a constant laboratory environment with an *ad libitum* food source (Schärer et al. 2005). In addition, antagonistic pleiotropy for sex-specific fitness and somatic maintenance has been previously documented in a quantitative genetic study of an Ascidian (Yund et al. 1997), which at least suggests that it is possible that selection on fitness via the female sex role could result in a correlated reduction in fitness via the male sex role.

### Effects on male-specific fitness

Given the changes we observed in sex-specific fitness in the female-selected lines, why was there no difference observed in the male-selected lines compared to the control lines? Sex-limited experimental evolution studies in *Drosophila* have generally detected a larger response to male-specific selection compared to female-specific selection (discussed in Abbott et al. 2020), so our results are somewhat surprising in this respect. There are at least two plausible explanations which are not mutually exclusive. Firstly, Nordén and Abbott (2017) showed that the additive genetic variance for female fitness was significantly higher than additive genetic variance for male fitness in the ancestral population that was used to set up the experimental evolution lines. The response in the male-selected lines could therefore have been constrained by the lower amount of additive genetic variation for male-specific fitness. Secondly, our experimental set-up is likely to be more efficient at selecting on female fitness than male fitness. *M. lignano* has been observed to occur at high population densities in the field (Ladurner et al. 2005), and seems to mate frequently both in the field and in the laboratory (Janicke et al. 2016). Our selection protocol restricted the number of mating partners to two for logistical reasons. This is a smaller mating group than the laboratory-adapted populations usually experience (normally ~100, Marie-Orleach et al. 2014), and likely smaller than those under natural conditions as well (Ladurner et al. 2005). Given that post-copulatory sexual selection is important for the male sex role in this species (Marie-Orleach et al. 2016), the number of mating partners in the experiment might have limited the scope for selection on sperm competitive ability, and thus restricted the opportunity for selection on male fitness.

Even though we did not find that the male-selected lines had changed relative to the control lines, we did find that relative testes area significantly predicted male-specific fitness on the line level. Similar patterns are well-established in separate-sexed species (reviewed in Lüpold et al. 2020), and previous results have shown that manipulation of group size results in changes in testis size and spermatogenesis in *M. lignano* (Janicke et al. 2013, Sekii et al. 2013, Giannakara et al. 2016, Vellnow et al. 2017, Nieuwenhuis and Aanen 2018). This result is therefore perhaps unsurprising, but at least suggests that our experimental evolution protocol does not seem to have unexpectedly altered this fundamental relationship.

### Conclusions and implications for sex chromosome evolution

Our main aim with this experiment was to attempt to mimic the first steps in a transition from hermaphroditism to separate sexes (Charlesworth and Charlesworth 1978, Bachtrog et al. 2014, Beukeboom and Perrin 2014) in the laboratory. We feel that the results presented here are proof of concept that this scenario is - at least to some extent - observable in real time. The first steps towards the evolution of a new sex chromosome system are thought to occur when a sterility mutation in one sex role becomes linked to one or more sex-specific fitness loci, leading to increased fitness in that sex (Beukeboom and Perrin 2014, Zemp et al. 2018) We do not know at this stage whether the changes in fitness we have observed are mainly controlled by loci linked to the GFP marker used in the selection protocol or not. However, we have shown that the sex-specific fitness changes thought to be associated with early sex chromosome evolution can be re-created in the laboratory, without directly manipulating sex allocation. Although it would also be possible to study the evolution of nascent sex chromosomes by inducing true sterility in one or the other sex role, the advantage with our experimental set-up is that we can design experiments to disentangle the sexual antagonism and sex allocation hypotheses. Work in this direction is ongoing.

According to the generally-accepted theory of sex chromosome evolution, the next step would be the evolution of linkage between sex-specific fitness loci and the sex-determining locus, and later recombination arrest around the sex-determining region (Charlesworth et al. 2005, Beukeboom and Perrin 2014). We would then expect to see that changes in gene expression or allele frequency are more likely to occur in loci that are physically linked to the GFP marker, although it is unclear whether such changes are likely to occur on a short enough time scale to be detectable over the course of the experiment. Still, we have shown that sex-specific selection in a hermaphrodite can help us to gain insight into the very earliest stages in sex chromosome evolution.

## Acknowledgements

We thank Lukas Schärer for important feedback when planning the study. We also thank Tammy Ho who helped with lab work during the fitness assay and Dita Vizoso for laboratory protocols and methods. This project was funded by Maja och Erik Lindqvists forskningsstiftelse, Nilsson-Ehle donationerna, the Crafoord Foundation 20120628 and 20140644, VR 2011-05679 and 2015-04680, and ERC-Stg-678148.

## Supplemental information

**Table S1:**
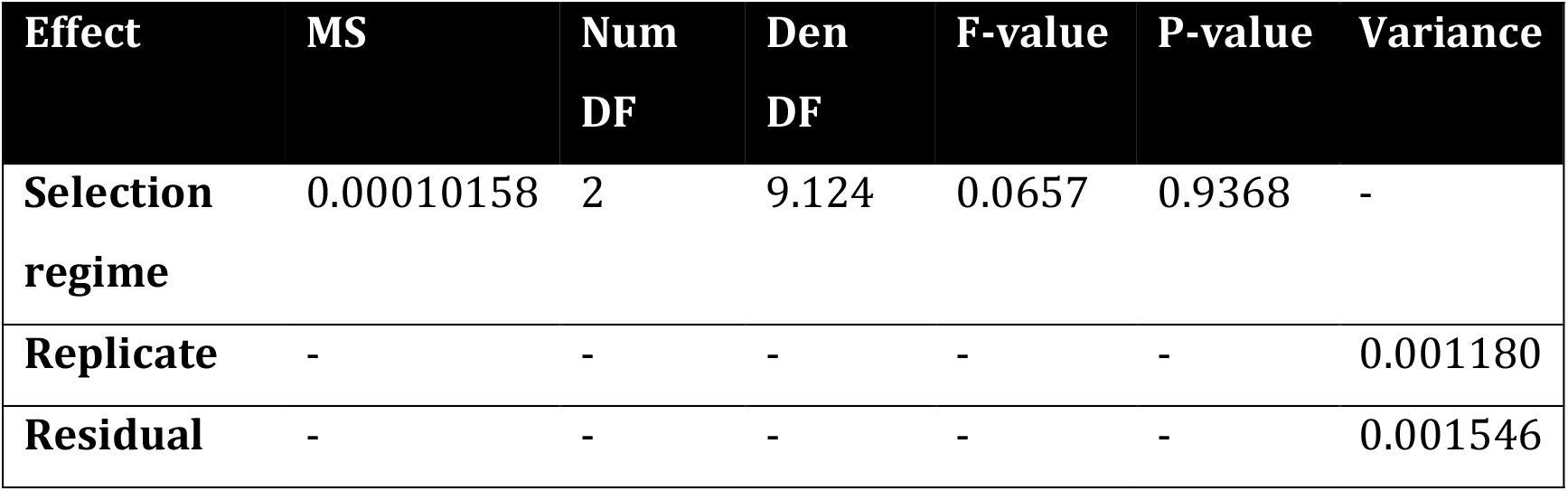
Results of the mixed model analysis of body area.

| Effect | MS | Num<br>DF | Den<br>DF | F-value | P-value | Variance |
| --- | --- | --- | --- | --- | --- | --- |
| **Selection regime** | 0.00010158 | 2 | 9.124 | 0.0657 | 0.9368 | - |
| **Replicate** | - | - | - | - | - | 0.001180 |
| **Residual** | - | - | - | - | - | 0.001546 |

**Table S2:** Results of the mixed model analysis of A) relative ovary area, and B) ovary area with body size as a covariate.

| Effect | MS | Num<br>DF | Den<br>DF | F-value | P-value | Variance |
| --- | --- | --- | --- | --- | --- | --- |
| **Selection regime** | $3.160 \times 10^{-5}$ | 2 | 8.243 | 0.4118 | 0.6753 | - |
| **Replicate** | - | - | - | - | - | $9.323 \times 10^{-6}$ |
| **Residual** | - | - | - | - | - | $7.672 \times 10^{-5}$ |

B)
| Effect | MS | Num<br>DF | Den DF | F-value | P-value | Variance |
| --- | --- | --- | --- | --- | --- | --- |
| **Body area** | $1.493 \times 10^{-4}$ | 1 | 77.011 | 50.93 | $4.57 \times 10^{-10}$ | - |
| **Selection regime** | $5.060 \times 10^{-7}$ | 2 | 8.024 | 0.1726 | 0.8445 | - |
| **Replicate** | - | - | - | - | - | $4.065 \times 10^{-7}$ |
| **Residual** | - | - | - | - | - | $2.930 \times 10^{-6}$ |

**Table S3:** Results of the mixed model analysis of A) relative testis area, and B) testis area with body size as a covariate. (The high denominator degrees of freedom in the analysis of testis area with body size as a covariate are a result of the fact that zero variance could be assigned to replicate population for this trait.)

| Effect | MS | Num<br>DF | Den<br>DF | F-value | P-value | Variance |
| --- | --- | --- | --- | --- | --- | --- |
| **Selection regime** | 2.654*10<sup>-5</sup> | 2 | 110 | 0.2512 | 0.7783 | - |
| **Replicate** | - | - | - | - | - | 0.0000 |
| **Residual** | - | - | - | - | - | 0.0001056 |

B)
| Effect | MS | Num<br>DF | Den<br>DF | F-value | P-value | Variance |
| --- | --- | --- | --- | --- | --- | --- |
| **Body area** | 2.761*10<sup>-4</sup> | 1 | 109 | 66.27 | 6.996*10<sup>-13</sup> | - |
| **Selection regime** | 1.340*10<sup>-7</sup> | 2 | 109 | 0.0322 | 0.9683 | - |
| **Replicate** | - | - | - | - | - | 0 |
| **Residual** | - | - | - | - | - | 4.166*10<sup>-6</sup> |

